# Monitoring of *Plasmodium falciparum* and *Plasmodium vivax* using microsatellite markers indicates limited changes in population structure after substantial transmission decline in Papua New Guinea

**DOI:** 10.1101/817320

**Authors:** Johanna Helena Kattenberg, Zahra Razook, Raksmei Keo, Cristian Koepfli, Charlie Jennison, Dulcie Lautu-Ninda, Abebe A. Fola, Maria Ome-Kaius, Céline Barnadas, Peter Siba, Ingrid Felger, James Kazura, Ivo Mueller, Leanne J. Robinson, Alyssa E. Barry

## Abstract

Monitoring the genetic structure of malaria parasite populations has been proposed as a novel and sensitive approach to quantify the impact of malaria control and elimination efforts. Here we describe the first population genetic analysis of sympatric *Plasmodium falciparum (Pf) and Plasmodium vivax (Pv)* populations following nationwide distribution of long-lasting insecticide treated nets (LLIN) in Papua New Guinea (PNG). Parasite isolates from serial cross-sectional studies pre-(2005-6) and post-LLIN (2010-2014) were genotyped using microsatellite markers. Despite parasite prevalence declining substantially in these communities (East Sepik: *Pf*=54.9-8.5%, *Pv*=35.7-5.6%, Madang: *Pf*=38.0-9.0%, *Pv*: 31.8-19.7%), genetically diverse and intermixing parasite populations remained. *P. falciparum* diversity declined modestly post-LLIN relative to pre-LLIN (East Sepik: *R*_s_ = 7.1-6.4, *H*_e_ = 0.77-0.71; Madang: *R*_s_= 8.2-6.1, *H*_e_ = 0.79-0.71). Unexpectedly, population structure present in pre-LLIN populations was lost post-LLIN, suggesting that more frequent human movement between provinces may have contributed to higher gene flow between provinces. *P. vivax* prevalence initially declined but increased again in one province, yet diversity remained high throughout the study period (East Sepik: *R*_s_=11.4-9.3, *H*_e_=0.83-0.80; Madang: *R*_s_=12.2-14.5, *H*_e_=0.85-0.88). Although genetic differentiation values increased between provinces over time, no significant population structure was observed at any time point. For both species, the emergence of clonal transmission and significant multilocus linkage disequilibrium (mLD) due to increased focal inbreeding post-LLIN was a strong indicator of impact on the parasite population using these markers. After eight years of intensive malaria control in PNG and substantial prevalence decline the impact on parasite population diversity and structure detectable by microsatellite genotyping was limited.

## INTRODUCTION

While half of the world’s population is still at risk of malaria infection, between 2010 and 2016, intensified malaria control efforts worldwide have reduced disease incidence by 18% and mortality by 32% (*World Malaria Report 2017*, 2017). Increased financial support and wide-scale deployment of malaria control interventions has enabled these reductions in transmission in many endemic regions. Nevertheless, the most recent WHO World Malaria Report (*World Malaria Report 2018*, 2018) reveals that progress has stalled since 2016, with malaria resurging in some areas, signalling that without sustained control pressure and new control strategies, these gains could be reversed. Current malaria control and elimination programs face many challenges, including the increasing heterogeneity of transmission as prevalence declines and the unknown origins of infections at very low transmission. In addition, changing government priorities heralds the need for more extensive surveillance and understanding of the impact of control efforts (malEra Consultative Group on Monitoring & Surveillance, 2011).

Genomic surveillance of parasite populations has emerged as a promising and high-resolution approach for malaria monitoring (Arnott, Barry, & Reeder, 2012; Barry, Waltmann, Koepfli, Barnadas, & Mueller, 2015; Dalmat, Naughton, Kwan-Gett, Slyker, & Stuckey, 2019; Koepfli & Mueller, 2017; malEra Consultative Group on Monitoring & Surveillance, 2011). These approaches go beyond traditional epidemiological measures of malaria disease burden and infection prevalence by identifying local transmission dynamics (e.g. endemic, epidemic, imported infections) and the connectivity (gene flow) between parasite populations in different endemic areas (Anderson et al., 2000; Fola et al., 2017; Noviyanti et al., 2015; Vardo-Zalik et al., 2013; Waltmann et al., 2018). Regional surveys reveal source and sink populations and parasite migration in a country and can help to predict whether and where targeted interventions would be effective and the spatial scale required (Auburn & Barry, 2017; Barry et al., 2015; Koepfli & Mueller, 2017). Fine-scale population genetic surveys also identify local drivers contributing to sustained transmission such as particular human social and economic interactions (Barry et al., 2015; Delgado-Ratto et al., 2016; Koepfli & Mueller, 2017). While parasite population genetics and genomics is becoming more popular and accessible, the impact on control programs has been limited, and to date few studies have systematically assessed the long-term impact of malaria control using these approaches (Batista, Barbosa, Da Silva Bastos, Viana, & Ferreira, 2015; Bei et al., 2018; Branch et al., 2011; Chenet, Taylor, Blair, Zuluaga, & Escalante, 2015; R. F. Daniels et al., 2015; Gatei et al., 2010; Gunawardena, Ferreira, Kapilananda, Wirth, & Karunaweera, 2014; Iwagami et al., 2012; Salgueiro, Vicente, Figueiredo, & Pinto, 2016; Vardo-Zalik et al., 2013). Moreover, it is not clear how long transmission needs to be disrupted, or to which extent prevalence should be reduced, before changes in parasite population structure can be seen.

Based on microsatellite genotyping, the genetic diversity and population structure of the most virulent malaria parasite, *P. falciparum*, is closely associated with transmission intensity in different endemic regions, and therefore parasite population diversity and structure varies greatly on a global and even on a local scale (Anderson et al., 2000; Chenet, Schneider, Villegas, & Escalante, 2012; Gatei et al., 2010; Noviyanti et al., 2015; Orjuela-Sanchez et al., 2013; Pava et al., 2017; Salgueiro et al., 2016; Schultz et al., 2010; Vardo-Zalik et al., 2013)).

*P. falciparum* populations in low transmission regions generally have low proportions of multiclonal infections and high levels of multilocus linkage disequilibrium (mLD) suggesting significant inbreeding and infrequent recombination (Anderson et al., 2000; Branch et al., 2011; Chenet et al., 2012; Noviyanti et al., 2015). Conversely, populations in high transmission areas are often characterized by a high proportion of multiclonal infections and low levels of mLD (Anderson et al., 2000; Gatei et al., 2010; Orjuela-Sanchez et al., 2013; Salgueiro et al., 2016; Schultz et al., 2010). For *P. vivax*, a less virulent but nevertheless significant human pathogen, the picture is different. The impact of relapse and other unique features of *P. vivax* biology (Olliaro et al., 2016) on underlying transmission dynamics is evidenced by higher genetic diversity and multiclonal infections, even at low transmission (Arnott et al., 2013; Barry et al., 2015; Batista et al., 2015; Chenet et al., 2012; Delgado-Ratto et al., 2016; Ferreira et al., 2007; Fola et al., 2017; Fola et al., 2018; Gunawardena et al., 2014; Hong et al., 2016; Iwagami et al., 2012; Jennison et al., 2015; Koepfli, Rodrigues, et al., 2015; Noviyanti et al., 2015; Orjuela-Sanchez et al., 2013; Pava et al., 2017; Waltmann et al., 2018). *P. vivax* population genetic analyses in the South West Pacific region, reveal a modest decline in diversity and increasing population structure with the eastward decline in transmission (Fola et al., 2017; Koepfli et al., 2013; Waltmann et al., 2018). However, a clear signature of significant mLD has been observed at low transmission in several studies suggesting increasingly focal transmission as malaria rates decline, and the increasing contribution of genetically similar relapses as new infections become rarer (Barry et al., 2015; Batista et al., 2015; Chenet et al., 2012; Delgado-Ratto et al., 2016; Ferreira et al., 2007; Imwong et al., 2007; Iwagami et al., 2012; Noviyanti et al., 2015; Orjuela-Sanchez et al., 2013). *P. vivax* has higher diversity compared to *P. falciparum* which is consistent with a longer association with humans (Gilabert et al., 2018; Hupalo et al., 2016; W. Liu et al., 2014; Loy et al., 2017; Neafsey et al., 2012), a higher effective transmission intensity (likely due to relapse) (Hofmann et al., 2017; Lin et al., 2010; Robinson et al., 2015), and suggests that *P. vivax* populations have not undergone recent population bottlenecks as observed for *P. falciparum* (Hupalo et al., 2016; Neafsey et al., 2012).

Furthermore, persistent *P. vivax* transmission in areas where *P. falciparum* has been near to- or completely eliminated shows *P. vivax* is more resilient to control efforts and thus may be less likely to show changes in parasite population structure (Barry et al., 2015; Cornejo & Escalante, 2006; Feachem et al., 2010; W. Liu et al., 2014; Neafsey et al., 2012; Oliveira-Ferreira et al., 2010; Waltmann et al., 2015). Few studies have investigated the population genetics of sympatric *P. vivax* and *P. falciparum* populations (Chenet et al., 2012; Jennison et al., 2015; Noviyanti et al., 2015; Orjuela-Sanchez et al., 2013; Pava et al., 2017) and none have systematically assessed the relative impact of intensified malaria control on these two species over an extended period. A better understanding of the impact of malaria control interventions on *P. falciparum* and *P. vivax* population structure is urgently required to capitalise on the potential of genomic surveillance for malaria control and elimination.

In the WHO Western Pacific Region (WPR), the malaria mortality rate declined by 58% over the period 2010–2015, however infection prevalence in Papua New Guinea (PNG) remains the highest in this region (and outside the African continent), contributing 81% of malaria cases and 86% of malaria deaths in 2017 in the region (*World Malaria Report 2017*, 2017; *World Malaria Report 2018*, 2018) primarily due to *P. falciparum* and *P. vivax* infections (Kattenberg, 2018; Koepfli et al., 2017; *World Malaria Report 2018*, 2018). In 2003, renewed political and financial will to strengthen malaria control worldwide resulted in a new national malaria control campaign to quickly achieve high levels of long-lasting insecticidal mosquito nets (LLIN) ownership and usage in PNG (Hetzel et al., 2012; Hetzel et al., 2014). Coverage with LLIN was low in most parts of the country before nationwide free distribution took place (2004-8 and 2009-2012) (Betuela et al., 2012; Genton et al., 1994; Hii et al., 2001). This resulted in a significant increase in ownership of bed nets across the country by 2010 (any type 80%; LLINs 65%) (Hetzel et al., 2012; Hetzel et al., 2014) and the average malaria incidence rate in sentinel sites dropped from 13/1,000 population to 2/1,000 (range 0.6-3.3/1000 post-LLIN) (Hetzel et al., 2016). Prevalence of malaria infections decreased significantly post-LLIN on a provincial level as determined by light microscopy (Hetzel et al., 2016), as well as with in-depth studies using molecular diagnostics on smaller geographic scales in the East Sepik (ESP) and Madang Provinces (MAD) on the hyperendemic north coast (Arnott et al., 2013; Barry et al., 2013; Kattenberg, 2018; Koepfli et al., 2017; Koepfli et al., 2015). *P. falciparum* prevalence by LDR-FMA or qPCR, dropped from 38% pre-LLIN to 12% post-LLIN and *P. vivax* prevalence decreased from 28% to 13% (Kattenberg, 2018; Koepfli et al., 2017). The prevalence reduction was stronger in East Sepik province, especially in the case of *P. vivax*, where parasites were detected by qPCR in only 6% of the participants in 2012/13 compared to 35.7% in 2005 (Kattenberg, 2018). Prevalence in different villages was heterogenous. For example, in ESP in 2012-13 0.6% to 61.9% of individuals per village were infected (any species, as detected by qPCR) (Kattenberg, 2018). In MAD, malaria prevalence post-LLIN was considerably lower than in 2006 (from 63% to 28% by qPCR), however, after an initial strong reduction in *P. vivax* prevalence was observed in 2010 (from 32% down to 13%, table 1), qPCR detected an increase of (mostly sub-microscopic) *P. vivax* infections to 20% in 2014 (Koepfli et al., 2017; Koepfli, Robinson, et al., 2015) (Figure 1). Reductions were more substantial for *P. falciparum* than for *P. vivax*, as has been seen in many co-endemic areas (Feachem et al., 2010).

**Figure 1.**
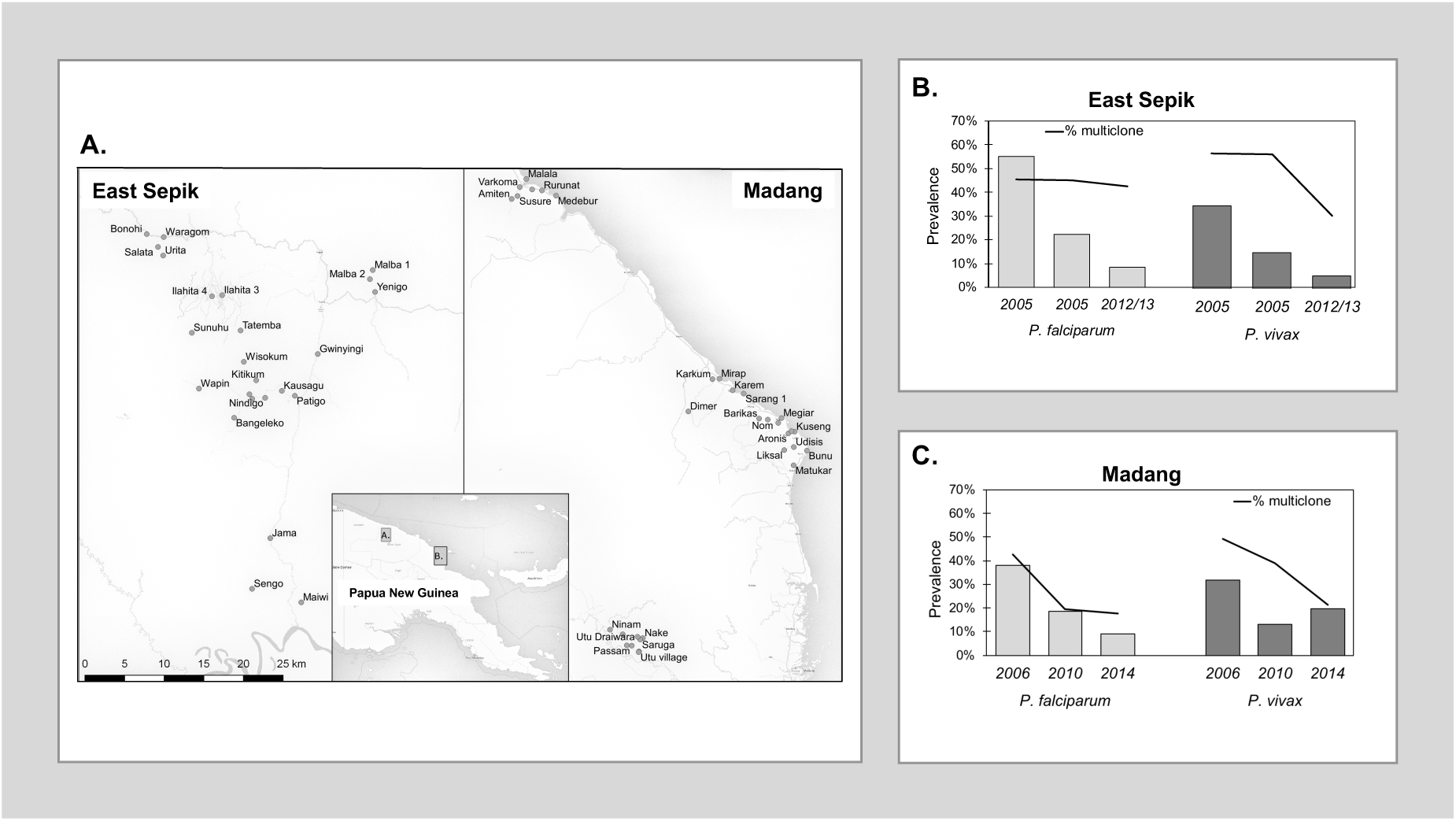
Map of the study areas and infection prevalence from 2005-2016. (A) Map of East Sepik and Madang study area villages and locations on the north coast of Papua New Guinea (inset) The graphs show the pre-LLIN (2005/6) and post-LLIN (2010-2014) molecular prevalence for (B) East Sepik and (C) Madang for both *P. falciparum* (light grey) and *P. vivax* (dark grey) and proportion of multiclonal infections (black line) (Data obtained from Arnott et al., 2013; Barry et al., 2013; Kattenberg, 2018; Koepfli et al., 2017; Koepfli et al., 2015; Mueller et al., 2009; Mueller et al., 2009; Senn et al., 2012).

Population genetic surveys using microsatellite markers were conducted in these areas of East Sepik and Madang Provinces before the intensification of malaria control locally (2005/2006). Higher genetic diversity and no population structure were observed for *P. vivax*, yet there was significant population structure of *P. falciparum* populations (Jennison et al., 2015; Koepfli et al., 2013; Schultz et al., 2010; Waltmann et al., 2018). Significant mLD was not observed for sub-populations of either species, confirming high levels of outcrossing and endemic transmission (Jennison et al., 2015). The impact of the decrease in prevalence post-LLIN on parasite population structure, however, remains unresolved. Therefore, we sought to characterise the post-LLIN diversity and population structure of sympatric *P. falciparum* and *P. vivax* populations using panels of well-validated neutral microsatellite markers (Anderson et al., 2000; Imwong et al., 2007; Karunaweera, Ferreira, Hartl, & Wirth, 2006). Microsatellite haplotypes were generated from *P. falciparum* and *P. vivax* samples collected after two rounds of mass LLIN distribution and compared to published data from isolates collected from the same two areas before the intensified malaria control program (Jennison et al., 2015; Schultz et al., 2010). The results show the impact of declining prevalence on parasite population diversity and structure and identify the critical parameters for monitoring these changes using microsatellite markers.

## MATERIALS AND METHODS

### Study sites and design

The studies were conducted in two Provinces on the highly endemic north coast region of Papua New Guinea (PNG) (Figure 1). In East Sepik Province, malaria decreased from a very high burden (73% of surveyed individuals infected in 2005 as measured by molecular detection (LDR-FMA (Mueller et al., 2009) to heterogeneous transmission (prevalence in villages ranging from 1% to 61%, median 6%, as measured by qPCR (Kattenberg, 2018)) after two rounds of LLIN distribution. An initial round of LLIN distribution was conducted between 2004 and 2009, followed by additional distributions in 2011/2012 and subsequently in 2014/15. In Madang province, however, malaria prevalence decreased from 63% to 28% by qPCR after the first round of LLIN distributions (Koepfli et al., 2017; Koepfli et al., 2015; Schultz et al., 2010). After the second LLIN distribution (2010-2014), *P. falciparum* continued to drop, however an increase in *P. vivax* prevalence was observed (from 13% to 20% by qPCR (Koepfli et al., 2017)). In Madang province, malaria prevalence was less heterogenous in the sampled villages than in East Sepik Province (Kattenberg, 2018; Koepfli et al., 2017; Koepfli et al., 2015).

Whole blood samples were collected from participants in cross-sectional studies conducted between 2005 and 2014 along the North Coast of PNG (Figure 1) (Mueller et al., 2009; Schultz et al., 2010). In Madang Province (MAD), the same three catchment areas were studied in 2006 (Schultz et al., 2010; Senn et al., 2012), 2010 (Koepfli et al., 2015) and 2014 (Koepfli et al., 2017). The study area included a selection of villages along a coastal stretch of 70km (Mugil and Malala regions), surrounded by coconut and cocoa plantations and subsistence gardens, and one area approximately 50 km inland (Utu). Here, the climate is tropical with a rainy season from December to April. In East Sepik Province (ESP), participants in the Wosera Catchment (ESP1) including fourteen villages were sampled in 2005 during the dry-season (August-September) (Jennison et al., 2015; Senn et al., 2012). A broader survey (ESP2) was conducted in April-May 2005 including five catchment areas that were re-visited in 2012-13 (Kattenberg, 2018; Mueller et al., 2009). The study areas in ESP consist of an area of over 160 km^2^ with low hills and riverine plains with a wet, tropical climate (Genton et al., 1995; Mueller et al., 2009). The natural vegetation is lowland hill forest that has mostly been replaced by re-growth following cultivation and wide grasslands on the plains near the Sepik River.

In all surveys, demographic and clinical information was collected, blood slides examined by expert microscopists and a blood sample collected in EDTA tubes for extraction of DNA. In the 2005 ESP studies, *Plasmodium* species were detected by Light Detection Reaction-Fluorescent Microspere Assay (LDR-FMA) (McNamara et al., 2006), whereas in all other studies quantitative PCR (qPCR) detection by TaqMan™ assay was used (Anna Rosanas-Urgell, 2010). To determine multiplicity of infection (MOI), *P. falciparum* positive samples were genotyped for *Pfmsp2* and *P. vivax* positive samples were genotyped with *Pvmsp1f3* and *MS16* (ESP1 and MAD 2006) or *Pvmsp1f3* and *MS2* (MAD 2010 & 2014 and ESP2 2005 & ESP 2012-13)), as previously described (Arnott et al., 2013; Kattenberg, 2018; Koepfli et al., 2017; Koepfli et al., 2015; Mueller et al., 2009; Schultz et al., 2010). The sample selection and genotyping procedures of the ESP1 2005 (Wosera) and MAD 2006 were as previously described (Arnott et al., 2013; Jennison et al., 2015; Schultz et al., 2010). For the other studies, samples with MOI of 1 were selected for further genotyping with the neutral microsatellite panels as described below. For the studies conducted after the large scale LLIN distribution (>2006) all monoclonal isolates (MOI=1) were included, but for the 2005 ESP2 population, a selection of samples was made for the analysis with the microsatellite panel (Table S1).

### Ethical approval

Written informed consent was obtained from all study participants or their parents or legal guardians. The study was approved by the PNG IMR Institutional Review Board (IRB#11/16) and the PNG Medical Research Advisory Committee (MRAC 11/21), National Institutes of Health, Division of Microbiology and Infectious Diseases (DMID Protocol #10-0035) and Walter and Eliza Hall Institute Human Research Ethics Committee (HREC #12/10).

### Genotyping procedures

For both species, a panel of 9-10 neutral microsatellite markers were amplified in the selected samples (Table S1) using a multiplex primary PCR followed by individual nested PCRs as previously described (Anderson et al., 2000; Jennison et al., 2015; Koepfli et al., 2013; Schultz et al., 2010). For *P. falciparum*, samples were genotyped at nine previously validated and commonly used, putatively neutral, microsatellite loci including *TA1, TAA60, Polya, ARA2, Pfg377, TAA87, PfPK2, TAA81* and *2490* (Anderson et al., 2000; Schultz et al., 2010). For *P. vivax*, 10 putatively neutral microsatellites were genotyped as previously described: *MS1, MS2, MS5, MS6, MS7, MS9, MS10, MS12, MS15*, and *MS20* (Jennison et al., 2015; Koepfli et al., 2013). All PCR products were sent to a commercial facility for fragment analysis on an ABI3730xl platform (Applied Biosystems) using the size standard LIZ500. Primers used were the same for all datasets (Jennison et al., 2015; Schultz et al., 2010).

### Analysis

The electropherograms were analysed with Genemapper V4.0 (Applied Biosystems) with the same peak calling strategy as described previously (Jennison et al., 2015; Schultz et al., 2010). To avoid artefacts, precautions were taken to ensure allele calling was consistent (Jennison et al., 2015), and carefully reconstructing dominant haplotypes as per previously described methods (Anderson, Su, Bockarie, Lagog, & Day, 1999; Jennison et al., 2015; Schultz et al., 2010). Briefly, this involved setting the minimum fluorescence to 500 Random Fluorescence Units (RFU) for all colours except the size standard. Stutter window was set to 3.5 for 3bp repeats and 4.5 for 4bp repeats. The stutter ratio was set to 0.4 for all markers. The stutter detection was only applied to shorter alleles, with longer alleles within the stutter window subject to the standard 30% cut-off threshold. Samples with low fluorescence were manually reanalysed with a minimum fluorescence of 100 RFU. For the Madang 2005 and Wosera 2006 *P. falciparum* data, previously published cleaned and rounded microsatellite allele repeat numbers for *P. falciparum* single clone infections (Schultz et al., 2010) were converted back to allele sizes using the known number of nucleotides/repeat, whereas for *P. vivax* the raw data (allele calls) was available (Jennison et al., 2015). These data were combined with the newly generated MS data from the other studies before binning the alleles using the TANDEM software (Matschiner & Salzburger, 2009). Allele frequencies of the entire dataset (incl. previously genotyped datasets) were investigated and outlying alleles (most likely caused by PCR artefacts) were removed. Samples with missing data at six (60%) or more MS loci were excluded from further analysis. We attempted to calibrate the *P. falciparum* data from pre-LLIN Madang 2006 and Wosera (ESP1 2005) by converting rounded repeat numbers back to allele sizes, binning together with the newly generated data and removing outliers. However, there was strong population structure when compared to the new dataset, indicating experimental differences despite the use of the same protocols. Thus, we excluded direct comparisons between old and new datasets for *P. falciparum*.

To conduct the population genetic analyses, allele frequencies and input files for the various population genetics programs were created using CONVERT version 1.31 (Glaubitz et al.,) and PGD Spider version 2.1.0.1 (Lischer & Excoffier, 2012). Genetic diversity was measured by calculating the number of alleles (*A*), Nei’s gene diversity (expected heterozygosity (*He*) (Nei, 1987)) and allelic richness (*Rs*) (Hurlbert, 1971) that corrects for sample size, using FSTAT version 2.9.3.2 (Goudet, 1995). Pairwise genetic differentiation was measured by calculating pairwise Jost’s *D* (Jost, 2008) and Weir and Cockerhams *F*_ST_ (Weir & Cockerham, 1984) values and 95% confidence intervals were estimated with 1000 bootstraps using the diveRsity package in R (Keenan, McGinnity, Cross, Crozier, & Prodöhl, 2013). In contrast to some earlier studies (Schultz et al., 2010), where haploid genotypes were coded as diploid genotypes, but homozygote at each locus, the data in this study was analysed using haploid datasets (as in Jennison *et al*. (Jennison et al., 2015)). As a measure of inbreeding in each population, multilocus linkage disequilibrium (mLD) was calculated using LIAN version 3.7, applying a Monte Carlo test with 100,000 re-sampling steps (Haubold & Hudson, 2000). In the LIAN analysis only samples with complete haplotypes were included. For both species the dominant and single haplotypes were compared within catchments to identify any significant linkage disequilibrium (mLD) using LIAN (Haubold & Hudson, 2000) and *F*_*ST*_ using FSTAT version 2.9.3.2 (Goudet, 1995).

Bottleneck (Piry, Luikart, & Cornuet) was used to test for an excess of heterozygosity brought about by the loss of rare alleles following a population bottleneck. A two-phase mutational (TPM) model of 70% stepwise and 30% non-stepwise mutations and run 1000 iterations was used. In addition, the allele frequency distribution for all loci was examined for a ‘mode shift’ in the distribution (Luikart, Allendorf, Cornuet, & Sherwin, 1998). To investigate parasite population genetic structure, the Bayesian clustering software, STRUCTURE version 2.3.4 (Pritchard, Stephens, & Donnelly, 2000) was used to investigate whether haplotypes for each species clustered according to geographical origin and/or within time periods. The analysis was run 20 times for K = 1 to 15 for 100,000 Monte Carlo Markov Chain (MCMC) iterations after a burn-in period of 10,000 using the admixture model and correlated allele frequencies. The second order rate of change of LnP[D], ΔK was calculated according to the method of Evanno et al. (Evanno, Regnaut, & Goudet, 2005) to determine the most likely K (most likely number of populations). CLUMPP version 1.1.2 (Jakobsson & Rosenberg, 2007) was used to facilitate the interpretation of population genetic results using files generated with STRUCTURE HARVESTER Web v0.6.94 (Earl & vonHoldt, 2012) and *Distruct 1.1* (Rosenberg, 2004) was used to visualize the structure plots with the data generated with CLUMPP. Statistical analysis of molecular epidemiological and population genetic parameters was done using non-parametric methods as indicated in the results-section using STATA v12.1 (StataCorp, USA). QGIS 2.14.12 (OpenSource Geospatial Foundation) was used to map the villages and households and maps were constructed using OpenStreetMap (contributors) background from the OpenLayers plugin.

## RESULTS

### Multiplicity of Infection

Multiplicity of infection (MOI), determined by genotyping highly polymorphic markers and counting the numbers of alleles in each infection, is a proxy measure of the intensity of transmission. MOI in all areas was lower in all areas post-LLIN distribution decreasing from 1.8 to 1.3 (p=0.0463 Mann Whitney U test) and 2.0 to 1.4 (p=0.0495 Mann Whitney U test), respectively for *P. falciparum* and *P. vivax*. This corresponds with a lower proportion of multiclonal infections, except for *P. falciparum* in ESP in 2012-13 (Figure 1 and Table S1). Despite an increase in PCR prevalence of *P. vivax* in Madang Province in 2014, few multiclonal infections were detected (Figure 1 and Table S1).

### Microsatellite haplotypes

We then genotyped parasite isolates at microsatellite loci to generate multilocus haplotypes for population genetic analyses. Multilocus haplotypes with at least five loci successfully genotyped (out of 9 for *P. falciparum* and 10 for *P. vivax*) were constructed for 860 *P. falciparum* samples (300 previously published (Jennison et al., 2015; Schultz et al., 2010)), and 755 *P. vivax* samples (202 previously published (Jennison et al., 2015; Schultz et al., 2010)) (Table S1). Despite having genotyped the samples that were identified as MOI=1 by *pfmsp2*, *pvmsp1f3* and *pvMS2/MS16* genotyping, 31% of *P. falciparum* samples and 49% of *P. vivax* samples had more than one allele for at least 1 microsatellite locus, suggesting multiple clone infection and the increased resolution of the microsatellite panel. From these we created dominant haplotypes (Schultz et al., 2010). No significant changes in multilocus Linkage Disequilibrium (mLD) were found when comparing single vs. all haplotypes combined within each study (Table S2). Low genetic differentiation was found between single and dominant haplotypes for *P. falciparum* in MAD2014 (*F*_ST_=0.063, p = 0.58), however, this can be explained by small sample size (n=9 dominant haplotypes). For *P. vivax*, low differentiation between single and dominant haplotypes in ESP 2012 (*F*_ST_ = 0.041, p = 0.33), was explained by a cluster of closely related haplotypes, all reconstructed from dominant alleles), which are described in more detail below. The fact that these related haplotypes were independently constructed from dominant haplotypes provides additional confidence in the allele-calling strategy. All other comparisons (within each province for at each time point) showed negligible genetic differentiation between single and dominant haplotypes. Therefore, the haplotypes were combined for further analysis.

### Reduction in *P. falciparum* but not *P. vivax* genetic diversity post-LLIN

Based on the microsatellite haplotypes (n=860), the genetic diversity of *P. falciparum* populations was modestly but significantly lower post-LLIN compared to the earlier time points for pre-LLIN populations (ESP1 and 2 2005 and MAD 2006). Mean heterozygosity for *P. falciparum* over all areas combined decreased significantly from 0.76±0.1 to 0.71±0.1 (Mann-Whitney U test p =0.036; Table S1) and allelic richness from 7.7±2.2 to 6.5±2.1 (Mann-Whitney U test p=0.014; Figure 2). These parameters also showed a small but insignificant decline for the provinces analysed individually (Figure 2, p>0.05). For *P. vivax*, overall genetic diversity remained high (post-LLIN *Rs=*12.5; *He*=0.85, Table S1), but slightly different results were seen in each province (Figure 2, Table S1). In Madang Province after the distribution of LLINs, *He* values slightly increased from 0.85±0.07 in 2006 to 0.88±0.04 in 2014 (p=0.3, Table S1), with high allelic richness (pre-LLIN *Rs* 12.2±4.0 vs post-LLIN 14.0±3.4; p=0.2). Whereas in East Sepik Province, *P. vivax* genetic diversity decreased but not significantly with *He* values of 0.83±0.09 to *He* 0.80±0.08 (2-sample t-test, p=0.48) and *Rs* values of 11.1±3.5 vs post-LLIN *Rs* 9.8±3.5 (2-sample t-test, p=0.33) (Table S1, Figure 2). No significant correlation was found between prevalence (by PCR) and heterozygosity, allelic richness, mean MOI, or proportion of multiple clone infections, for either species (data not shown).

**Figure 2.**
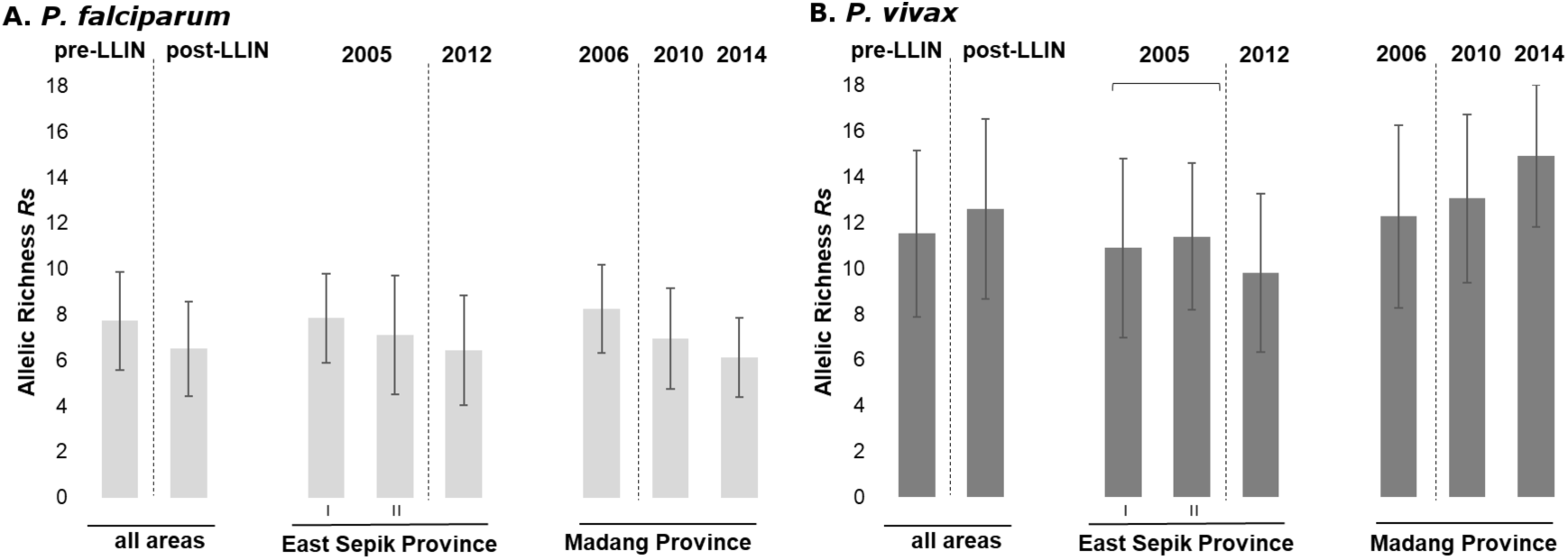
Changing diversity of *P. falciparum* and *P. vivax* populations over an intensifying malaria control period (2005-2014). Allelic Richness (*Rs*) in *P. falciparum* (A) (n= 860) and *P. vivax* (B) (n=755) populations pre-(≤2006) and post-LLIN (≥2010) mass-distributions. Error bars indicate standard deviations.

### Significant multilocus linkage disequilibrium (mLD) for both *P. falciparum* and *P. vivax* post-LLIN

For *P. falciparum*, matching haplotypes (allowing missing loci) were seen in all post-LLIN datasets and the pre-LLIN ESP2 2005 dataset. However, for *P. vivax*, matching haplotypes (allowing missing loci) were rarely seen and only in post-LLIN data sets. Among the 332 complete *P. falciparum* multilocus haplotypes (9-loci successfully genotyped) from all study sites, 16 repeated haplotypes were found, with 11 haplotypes represented two times, three represented three times, and two represented four times. Clonal haplotypes were always found within the same year and province, and in all cases except one in the same catchment area, but not always in the same village (7/16 haplotypes found in neighbouring villages). In ESP2 2005, one clonal haplotype was found in two villages (Yenigo and Sengo, Figure 1) from different catchment areas, roughly 40km apart. Among the 303 complete *P. vivax* multilocus haplotypes (10-loci), two haplotypes were repeated, with one haplotype represented two times (in two different villages in ESP 2012), and one represented four times (three in one village, one in a neighbouring village in MAD 2010).

To investigate whether inbreeding was present in these populations (Smith, Smith, O’Rourke, & Spratt, 1993), non-random associations among the microsatellite loci (mLD) were calculated for all complete and unique haplotypes. Whilst mLD was absent from most pre-LLIN populations, significant mLD was observed in *P. falciparum* and *P. vivax* infections post-LLIN (Figure 3). In ESP 2012-13, mLD was high with unique infections indicating the circulation of closely related haplotypes in the population, suggesting near clonal transmission (low recombination between diverse clones and high levels of inbreeding) or the presence of high proportions of meiotic siblings among isolates (Bright et al., 2014; Smith et al., 1993), as observed in the village of Sunuhu (Table S3). Low, but significant mLD was found for *P. falciparum* in Madang in 2006 (Figure 3), however this population was structured (Schultz et al., 2010), resulting in a phenomenon called the Wahlund effect, confirmed by the fact that linkage equilibrium was restored when mLD was analysed separately for subpopulations (Jennison et al., 2015; Wahlund, 1928) (Table S4). In post-LLIN Madang, the observed mLD for *P. falciparum* is not the result of subpopulation structure, as significant mLD remained in the subpopulations (Table S4). In Mugil area in 2014, significant mLD for *P. falciparum* remains due to the circulation of a few very closely related haplotypes in the Megiar village (Table S4, Dataset 1).

**Figure 3.**
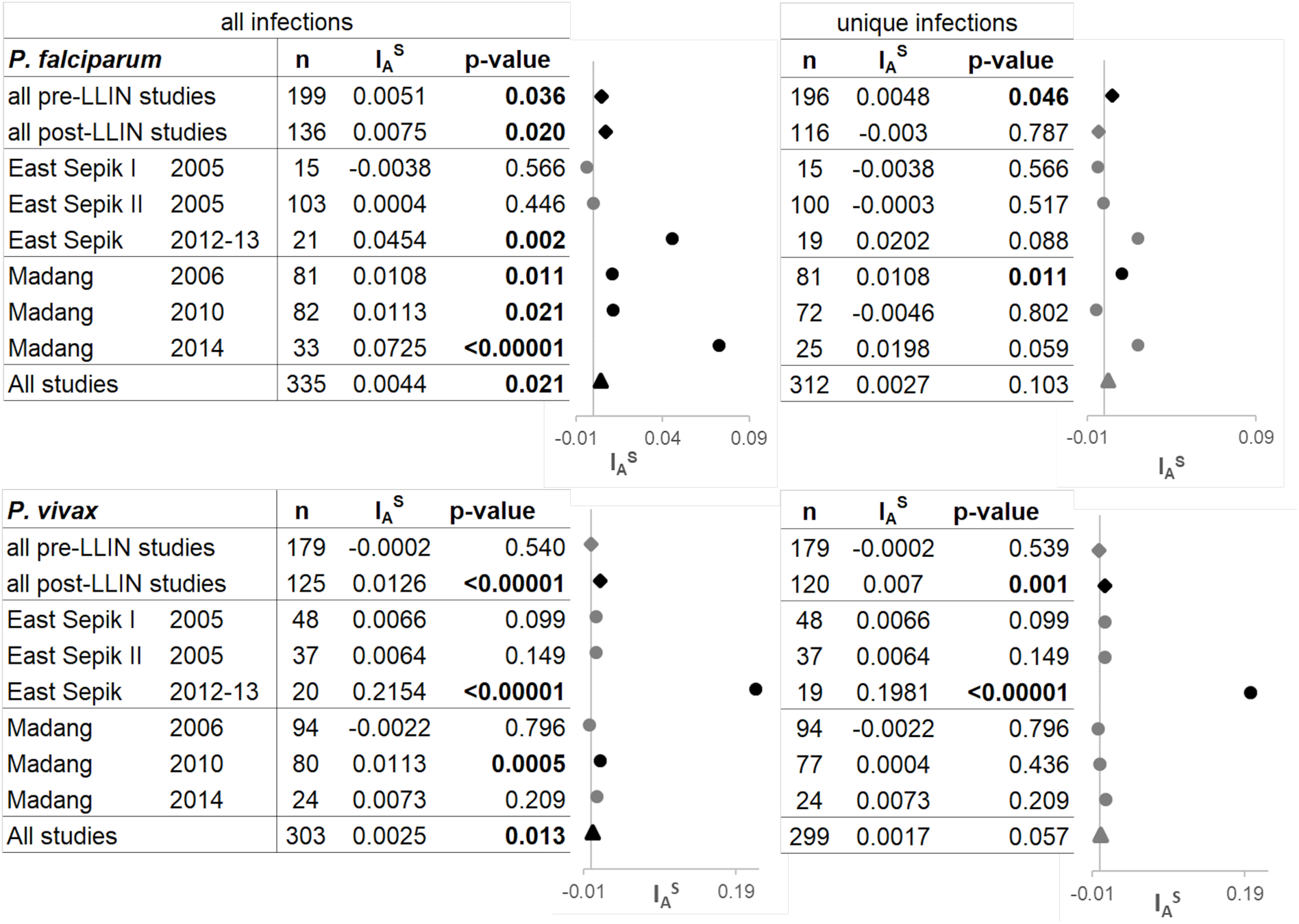
Estimates of multi-locus linkage disequilibrium (mLD) for *plasmodium* populations pre- and post-LLIN mass-distributions. The standardized index of association I_A_^S^) is plotted with black dots representing values with significant linkage disequilibrium; n = number of complete haplotypes. Single clone infection mLD in shown in Table S2.

### Population Bottleneck for *P. falciparum* but not *P. vivax* post-LLIN

Bottleneck analysis was performed using a two-phase mutational model (TPM) and testing for heterozygosity excess with a 2-tailed Wilcoxon sign rank test (see Materials and Methods). Significant heterozygosity excess was observed for *P. falciparum* in ESPII 2005 (but not in ESPI 2005), MAD 2006 and 2010 populations (p=0.020, p=0.049 and p= 0.010, respectively). However, for *P. vivax s*ignificant heterozygosity excess was only observed in a single pre-LLIN population, Wosera 2005 (p=0.042, these samples were collected in the relatively dry season) and not in any of the other *P. vivax* populations. Despite finding significant heterozygosity excess, a mode-shifted distribution of allele frequencies (as is frequently observed in bottlenecked populations) was not observed in any of the time points and provinces.

### Contrasting and dynamic patterns of population structure for both *P. falciparum* and *P. vivax*

As previously described, for *P. falciparum*, low to moderate genetic differentiation was seen between the Wosera 2005 (ESP1) and Madang 2006 studies (Jennison et al., 2015; Schultz et al., 2010) (Figure 4). Comparisons were not done between *Pf* 2005/6 and other *Pf* populations as it was not possible to calibrate data through combining allele calls before binning (see Materials and Methods). Post-LLIN there remains low to moderate genetic differentiation between ESP (2012-2013) and Madang (2014) (Figure 4). However, there was little genetic differentiation of East Sepik *P. falciparum* populations pre-LLIN (ESP2 2005) compared to post-LLIN (2012-13), nor between Madang 2010 and 2014 populations (Figure 4). For *P. vivax*, a different pattern can be seen, with low genetic differentiation between provinces pre-LLIN, which increases in the post LLIN-studies (Figure 4). Similar to *P. falciparum*, within province genetic differentiation between the different time points does not increase post-LLIN.

**Figure 4.**
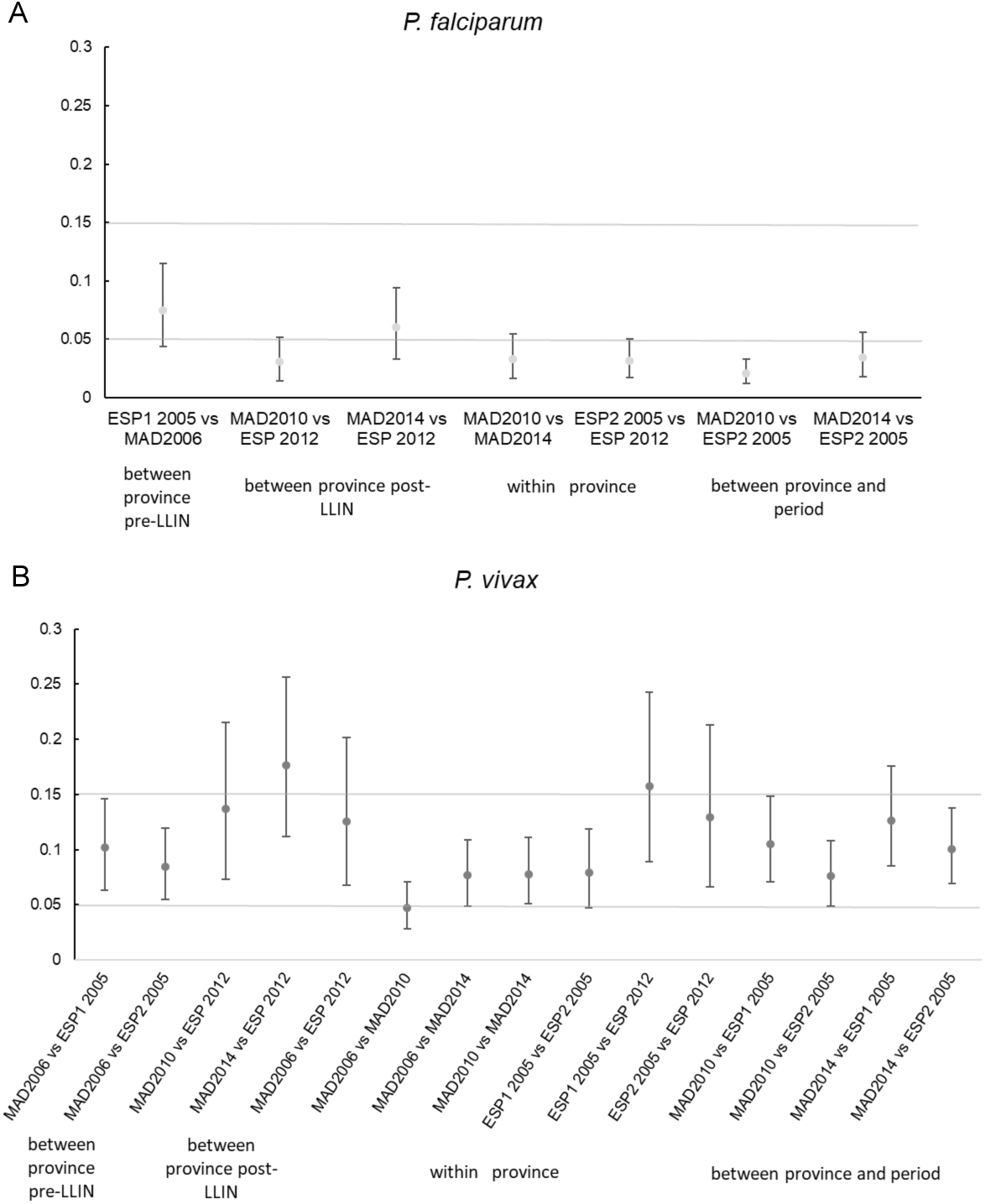
Genetic differentiation estimates among *P. falciparum* and *P. vivax* populations pre- and post-LLIN mass-distributions. Pairwise Jost’s D values for (A) *P. falciparum* and (B) *P. vivax.* Pairwise Jost’s D values and 95% confidence intervals were estimated with 1000 bootstraps using the diveRsity package in R. Pairwise F_ST_ values (Weir and Cockerham) are shown in figure S1.

Population genetic structure was further investigated by Bayesian cluster analysis using STRUCTURE (Pritchard et al., 2000). Haplotypes or populations with ancestry in more than one cluster are considered admixed and indicates that substantial gene flow occurs between defined geographic areas. Our analysis identified three *P. vivax* and three *P. falciparum* clusters (Figure 5, S7, S8 and S9). The clustering patterns show that the *P. falciparum* populations in later years are more mixed than the populations of 2005/2006, where populations clustered according to geographical locations including amongst the three catchment areas within Madang Province (Figure 5) (Schultz et al., 2010). On the contrary, *P. vivax* populations were very diverse and displayed little population structure in all time points, despite the increase in differentiation between ESP and MAD post-LLIN populations (Figures 4 and 5).

**Figure 5.**
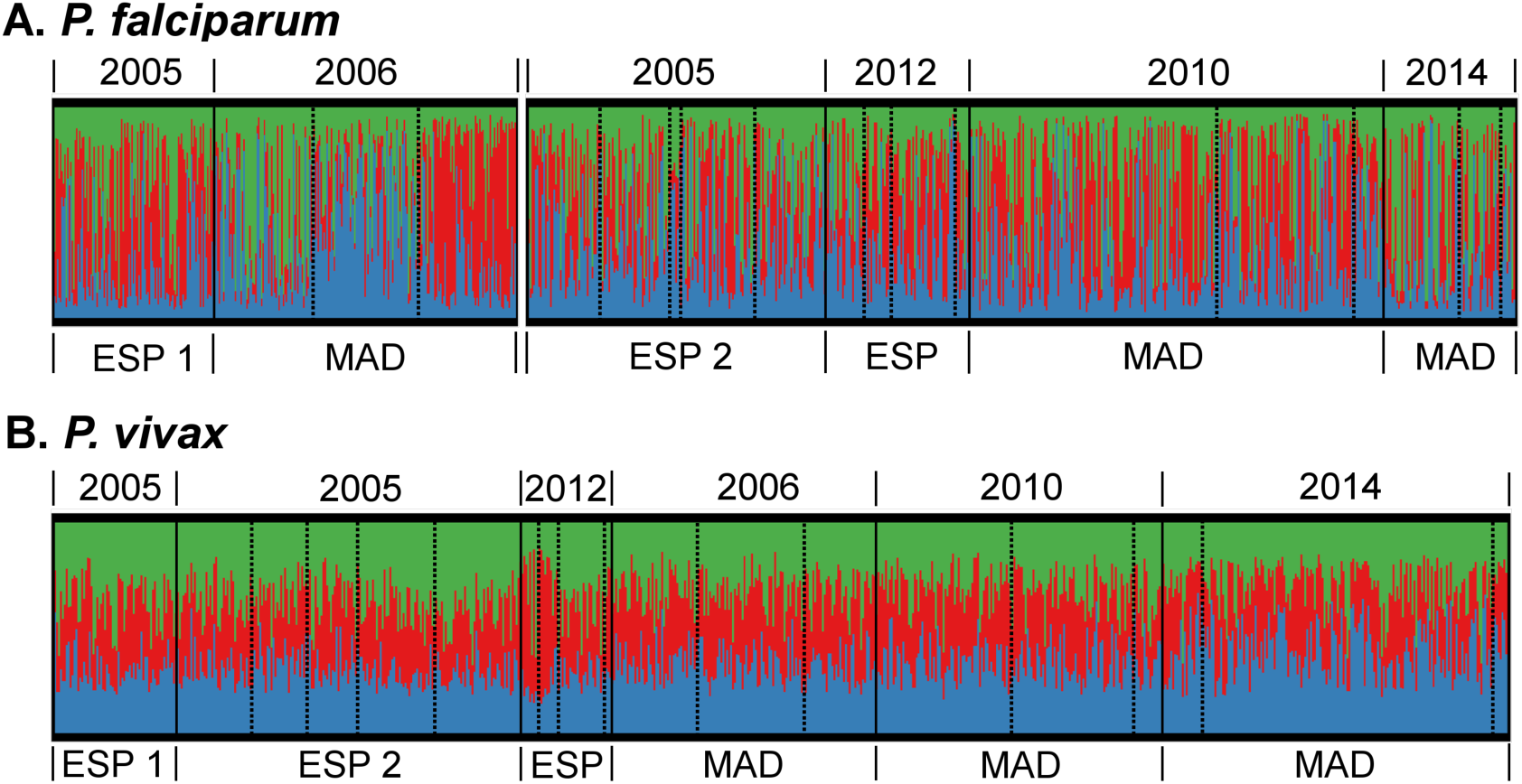
Bayesian cluster analysis of *P. falciparum* (A) and *P. vivax* (B) of pre-(2005/6) and post-LLIN studies (2010-2014). Individual samples are sorted by province and year (solid lines), catchment area (dashed line) and cluster membership (colour). Madang catchments are organised as Malala, Mugil, Utu and ESP2 and 2012-13 as Brukham, Burui, Ilahita, Ulupu, and Wombisa (no infections in 2012). As identified in the genetic differentiation analysis (Jost’s D, Figure 4), there was moderate differentiation for *P. falciparum* between the ESP1 (Wosera area) 2005 and Madang 2006 versus the other studies (only one province included pre-LLIN), which is believed to be for a large part caused by experimental- and data analysis differences. Therefore, these *P. falciparum* studies were grouped separately for the population structure analysis, in order to avoid artificial changes in ancestry over time.

For *P. vivax*, as we observed significant mLD post-LLIN in ESP, local phylogenetic analysis was conducted. This supports focal transmission as shown by the clustering of haplotypes from the same village: Sunuhu (Figure 6). Interestingly, this village had the highest prevalence in the region in the 2012-13 survey (36% infected with *P. vivax* by qPCR compared to 0.5-9% in other villages (Kattenberg, 2018). In Sunuhu, clonal and closely related haplotypes (≤2 unmatched alleles) were observed in 48% (11/23) of the haplotypes from that village (see supplementary file 1). The 11 closely related haplotypes were observed throughout the village, were not clustered in neighbouring households, and were not associated with participant characteristics (p>0.05), such as age and sex (Table S3).

**Figure 6.**
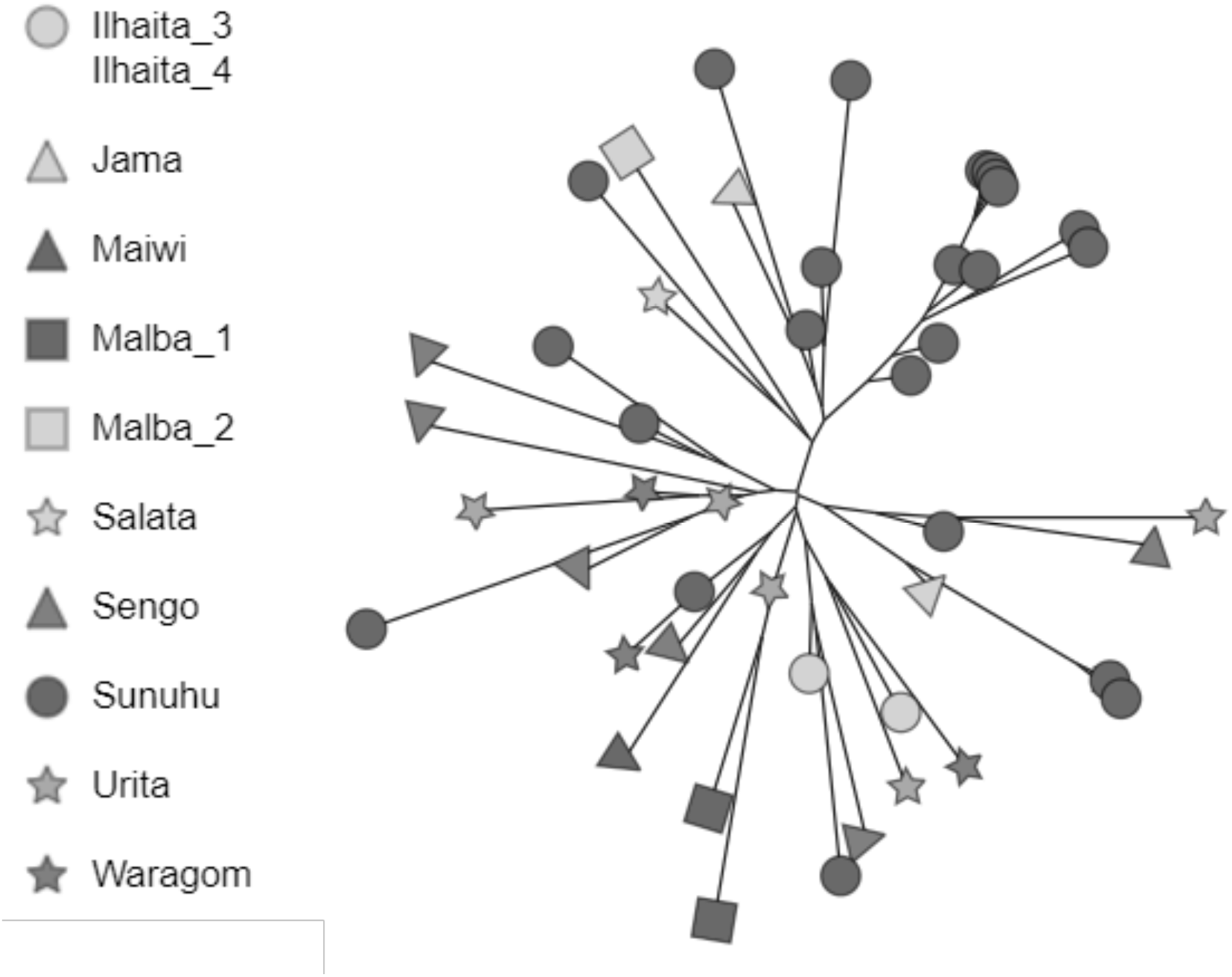
Unrooted neighbour-joining tree of *P. vivax* isolates in East Sepik province in 2012-13. Relatedness among haplotypes was defined by calculating the pairwise distance and neighbour-joining tree using the APE package in R and the tree was visualized using Phylocanvas through microreact.org (Argimón et al., 2016; “Microreact,”). Colours and shapes indicate the village where the isolates were collected. Very closely related isolates are observed in the village of Sunuhu (dark grey circles).

## DISCUSSION

Despite large reductions in parasite prevalence from very high to low/moderate levels in PNG following the nationwide rollout of LLIN (Kattenberg, 2018; Koepfli et al., 2017; Koepfli, Robinson, et al., 2015), we show that based on haplotypes of nine (*P. falciparum*) or ten (*P. vivax*) microsatellite alleles, parasite populations were minimally impacted and remain diverse and unstructured. *P. falciparum* diversity showed a minor decrease, though remained high relative to other malaria endemic areas outside Africa (Anderson et al., 2000; Branch et al., 2011; Chenet et al., 2012; Chenet et al., 2015; dalla Martha, Tada, Ferreira, da Silva, & Wunderlich, 2007; Noviyanti et al., 2015; Orjuela-Sanchez et al., 2009; Orjuela-Sanchez et al., 2013; Pava et al., 2017; Susomboon et al., 2008), whereas *P. vivax* diversity remained high throughout the study period. Surprisingly, *P. falciparum* populations that were structured pre-LLIN (2005, 2006) (Jennison et al., 2015; Schultz et al., 2010), were unstructured post-LLIN (2010, 2014), although focal transmission of clonal haplotypes and inbreeding was detected. For *P. vivax*, there was also no evidence of population structure after LLIN, however increasing pairwise genetic differentiation within and between provinces was observed and clonal transmission and inbreeding had emerged in at least one village.

For *P. falciparum*, transmission has previously been associated with decreasing diversity with increasing multilocus LD and population structure (Anderson et al., 2000; Chenet et al., 2012; Gatei et al., 2010; Noviyanti et al., 2015; Orjuela-Sanchez et al., 2013; Pava et al., 2017; Salgueiro et al., 2016; Schultz et al., 2010; Vardo-Zalik et al., 2013). In the East Sepik and Madang Provinces of PNG, *P. falciparum* infections decreased to very low prevalence in the human population. However, this resulted in only a minor decrease in parasite diversity and did not increase parasite population structure within (East Sepik only) or between provinces, indicating a large amount of gene flow between the sampled parasite populations. However, 17.7-42.4% of *P. falciparum* infections were multiclonal and may contribute to outcrossing and recombination during mosquito transmission [25, 73]. In addition, the lack of population structure indicates that high gene flow due to human migration may maintain this high diversity. For *P. falciparum*, the population structure observed prior to intensified control was thought to be the result of limited human migration or population bottlenecks in the past (Anderson et al., 2000; Jennison et al., 2015; Schultz et al., 2010; Tessema et al., 2015). PNG is home to extraordinary human linguistic and cultural diversity and extreme topographical features that have limited past human movement (I. Mueller, Bockarie, Alpers, & Smith, 2003; Riley, 1983). The two provinces included in this study are separated by roughly 300 hundred kilometres of mountainous and swampy terrain, are not connected by road and have substantial topographical and ecological differences (I. Mueller et al., 2003). In addition, changes in first-line treatment policies, for example the introduction of sulphadoxine/pyrimethamine in the early 2000’s and arthemeter-lumefantrine in 2008, might have played a role in shaping parasite population structure (Mu et al., 2005). CQ and/or SP resistance (near fixation of resistant *pfmdr1* and *pfdhps* resistant alleles were observed in the same areas (Barnadas et al., 2015; Koleala et al., 2015; Mita et al., 2018)) may have contributed to the observed bottleneck and mLD in pre-LLIN *P. falciparum* populations, with consequent reductions in effective population size, while outcrossing due to high transmission preserved within-population genetic diversity as the resistance mutation spread, as reported in another area (Mita et al., 2018). In later years, human mobility may have broken down *P. falciparum* population structure since large numbers of migrants from East Sepik have moved into Madang in the last decade seeking employment opportunities. Human mobility would have also maintained gene flow between *P. vivax* populations (Fola et al., 2018).

In contrast to *P. falciparum*, *P. vivax* diversity has previously been reported as high in both low and high transmission areas (Delgado-Ratto et al., 2016; Ferreira et al., 2007; Gunawardena et al., 2014; Hong et al., 2016; Noviyanti et al., 2015; Pava et al., 2017) or showing a modest decrease in diversity in low compared to high transmission areas (Fola et al., 2018; Iwagami et al., 2012; Waltmann et al., 2018), although clonal outbreaks and strong inbreeding can be observed in very low transmission areas (Abdullah et al., 2013; Batista et al., 2015; Fola et al., 2018; Y. Liu et al., 2014; Orjuela-Sanchez et al., 2013). Population structure patterns are also highly variable in different endemic areas (Abdullah et al., 2013; Delgado-Ratto et al., 2014; Ferreira et al., 2007; Hong et al., 2016; Koepfli, Rodrigues, et al., 2015; Koepfli et al., 2013; Pava et al., 2017). For the East Sepik *P. vivax* population, post-LLIN, significant LD was observed resulting from very closely related parasite strains circulating in a residual pocket of relatively high transmission within a single village. This suggests considerably reduced gene flow and inbreeding of parasites in that village, masked by relatively high overall genetic diversity and lack of evidence of a bottleneck at the Provincial level. This paradoxical signature of significant mLD with extensive population diversity and a considerable proportion of multiple clone infections of *P. vivax* appears to be a hallmark of lower transmission areas (Barry et al., 2015; Batista et al., 2015; Chenet et al., 2012; Delgado-Ratto et al., 2016; Delgado-Ratto et al., 2014; Ferreira et al., 2007; Hong et al., 2016; Noviyanti et al., 2015; Orjuela-Sanchez et al., 2013). Similar to *P. falciparum* populations though, there was a correlation between mLD and prevalence of infection for *P. vivax*. This shows that LD among microsatellite haplotypes is an earlier indicator of reduced transmission than genetic diversity and population structure (for both species).

Multilocus LD detected in post-LLIN *P. vivax* populations was explained by both focal epidemic expansions and inbreeding, as similarly observed in other studies in Peru, Vietnam, and Papua Indonesia (Delgado-Ratto et al., 2014; Hong et al., 2016; Noviyanti et al., 2015). The explanation for this observation will most likely lie in unique *P. vivax* characteristics related to hypnozoites, relapses and transmission dynamics (Abdullah et al., 2013; Bright et al., 2014; Chen, Auliff, Rieckmann, Gatton, & Cheng, 2007; Delgado-Ratto et al., 2014; Ferreira et al., 2007; Fola et al., 2018; Iwagami et al., 2012; White, 2011). At high transmission levels, as in pre-LLIN studies in PNG, which are characterised by high prevalence and high MOI, these clusters of similar haplotypes might also occur, but don’t impact multilocus LD and are obscured due to sampling limitations and the large number of different haplogroups circulating at the same time. As transmission declines, infections have fewer clones and the diversity of the hypnozoite reservoir decreases, resulting in fewer circulating haplogroups, lower levels of recombination between distinct genomes and more frequent clonal transmission, resulting in the observed LD as in this and other studies (Barry et al., 2015; Batista et al., 2015; Chenet et al., 2012; Delgado-Ratto et al., 2016; Delgado-Ratto et al., 2014; Ferreira et al., 2007; Hong et al., 2016; Noviyanti et al., 2015; Orjuela-Sanchez et al., 2013). These signatures are thus more likely to be seen in a population with sustained low transmission such as the East Sepik Province.

Considerable variance in the impact of the LLIN program was observed in the two provinces. In Madang, whilst *P. falciparum* rates steadily declined over the entire study period, there was a resurgence of submicroscopic *P. vivax* infections in 2014 (Koepfli et al., 2017). The lack of mLD and population structure suggests that this is not due to an outbreak, but more likely the residual *P. vivax* population was able to gain a foothold once again despite the intensive control efforts. In addition, an increase in the prevalence of *pvdhfr* triple and quadruple mutants, related with SP resistance, were observed in Madang province in 2010 (Barnadas et al., 2015), and a high proportion of resistant parasites could be a possible explanation for the higher infection prevalence by 2014. Different studies have shown that selective pressure of drugs such as chloroquine (CQ) and/or sulfadoxine-pyrimethamine (SP) have had an impact on shaping worldwide *P. vivax* populations in recent history (Hupalo et al., 2016; Pearson et al., 2016). However, a population bottleneck as seen in *P. falciparum* populations (Mita et al., 2018) did not occur in *P. vivax* populations of PNG. Malaria control had a greater impact on *P. vivax* prevalence in East Sepik and population structure was observed in one village post-LLIN. In this region, malaria transmission is heterogeneous between villages. Besides the national malaria control program, other initiatives were also distributing LLINs in East Sepik Province prior to the nationwide distribution (Hetzel et al., 2012; Hetzel et al., 2014; Hetzel et al., 2016) suggesting that longer term sustained control efforts have been effective.

This study has some limitations related to sampling and the genotyping approach used. The samples were collected in serial cross-sectional surveys over a period of malaria control initiated at different times in the two provinces. Several years of sustained control pressure in East Sepik might explain why, despite substantial prevalence decline in Madang Province by 2010 we did not observe any signature of changing population structure. Whilst, in East Sepik 2012, a cluster of closely related parasites was observed in one village suggesting more focal transmission than previous years. The microsatellite panels were selected as these have been the gold standard genotyping tool for large-scale malaria population genetic studies for many years (Anderson et al., 1999; Imwong et al., 2006; Karunaweera et al., 2006). However, they are limited in number (less than one per chromosome), rapidly evolving and prone to error. Although these markers are extremely useful for measuring parasite population structure on local and global scales (Auburn & Barry, 2017; Barry et al., 2015; Koepfli & Mueller, 2017; Sutton, 2013), they are not ideal for cross-study comparisons due to the difficultly in calibrating alleles from different data sources. The lack of raw data from the previously published *P. falciparum* study (Schultz et al., 2010), prevented the direct comparison of haplotypes and thus the analysis of population structure between the earlier *P. falciparum* time points for the East Sepik II (Wosera) and Madang populations. Furthermore, dominant haplotypes derived from multiple clone infections can be reconstructed incorrectly, thus inflating diversity values (Messerli, Hofmann, Beck, & Felger, 2017). However, only haplotype-based analyses such as multilocus LD and phylogenetic analysis were vulnerable to these possible artefacts. All other analyses were conducted using mean values across markers or allele frequencies and thus limit the impact of such errors. Moreover, the dataset included a large proportion of single infection haplotypes in all populations, and the detected clones included dominant haplotypes suggesting that allele calling was reliable. Finally, the highly polymorphic and rapidly evolving nature of microsatellite markers (Ellegren, 2004) may limit the resolution of the population genetic parameters such as population level diversity and population structure in high transmission areas (Branch et al., 2011). This may both lead to false assignment of unrelated parasites (e.g. from different provinces) as related, and may also reduce the detection of truly related parasites (identical by descent), both of which would indicate high gene flow and a lack of population structure. Other more stable markers, such as single nucleotide polymorphisms genotyped through barcoding (Baniecki et al., 2015; Daniels et al., 2008) or whole genome sequencing (Hupalo et al., 2016; Miotto et al., 2013; Mu et al., 2005; Pearson et al., 2016; Volkman et al., 2007) may be more sensitive to detect changes in parasite population structure. The high parasite diversity and lack of population structure are consistent with both species maintaining a large and evolutionarily fit population despite significant prevalence decline. However, the emergence of significant mLD indicates there is evidence of focal interrupted transmission and suggests that this parameter may be a useful marker for control impact and early changes in parasite population structure. *P. falciparum* population structure was lost even despite drastically declining transmission suggesting parallel changes in human behaviour with more frequent travel and immigration may have contributed to the patterns observed. These results were in contrast to our expectations of decreasing diversity and increasing population structure (Jennison et al., 2015) and show that long term sustained control efforts need to be maintained to observe significant increase in population structure at least as measured by the microsatellite panels used in this study. The PNG national malaria control program has made a huge impact on the malaria burden through the substantial reductions in infections circulating in the community (Hetzel et al., 2016; Kattenberg, 2018; Koepfli et al., 2017), however it appears that this has not been sustained long enough for the underlying parasite population to fragment. Finally, this study demonstrates how parasite population genetics can inform whether malaria intervention has reduced and fragmented the parasite population or if significantly more control effort will be required.

## ACKNOWLEDGEMENTS

We are grateful to the people who participated in the study, and for the support and collaboration of the participating communities. We appreciate the efforts of the laboratory, field team members and support staff from the Papua New Guinea Institute of Medical Research for their involvement in the included studies. This research was supported by National Health and Medical Research Council (NHMRC) of Australia Project Grants (1010069 and 10027108), the International Centres of Excellence in Malaria Research (ICEMR) for the South West Pacific, NIH U19 AI089686 and a Bill & Melinda Gates Foundation Grant (TransEpi Consortium). IM and LR are supported by NHMRC Research Fellowships.

## Data accessibility

**Dataset 1.** Microsatellite genotypes will be deposited in Dryad (https://datadryad.org)

## Author Contributions

AEB, IM and JK identified the research problem and designed the research project, MO-K, PS, IF, M, JK and LJR co-ordinated the field studies and molecular epidemiology, CK, DL-N, CB processed samples and performed molecular assays, JHK, ZR, RK and AAF conducted the microsatellite genotyping and analysed data, CJ and CK contributed data, JHK and AEB wrote the paper.

## SUPPORTING INFORMATION

**Table S1.** Genetic diversity of *P. falciparum* and *P. vivax* populations of Papua New Guinea over a period of intensifying malaria control (2005-2014).

| *P. falciparum* |  |  |  |  |  |  |  |  |  |  |  |  |  |  |
| --- | --- | --- | --- | --- | --- | --- | --- | --- | --- | --- | --- | --- | --- | --- |
|  | year | Province | N* | Prevalence* | mean<br>MOI* | %<br>multiclone<br>infections** | n<sub>s</sub> | n<sub>g</sub> | A±SD |  | Rs±SD |  | He±SD |  |
| pre-LLIN | 2005 | East Sepik 1 | 1077 | 22.3% | 1.7 | 45.0% | 113 | 102 | 9.4 | 2.8 | 7.8 | 2.0 | 0.77 | 0.13 |
|  | 2005 | East Sepik 2 | 2527 | 54.9% | 1.8 | 45.4% | 186 | 176 | 8.7 | 3.2 | 7.1 | 2.6 | 0.71 | 0.16 |
| post-LLIN | 2012-13 | East Sepik | 2486 | 8.5% | 1.6 | 42.4% | 95 | 79 | 7.4 | 3.2 | 6.4 | 2.4 | 0.71 | 0.12 |
| pre-LLIN | 2006 | Madang | 1282 | 38.0% | 1.9 | 42.7% | 207 | 198 | 10.8 | 2.9 | 8.2 | 1.9 | 0.79 | 0.07 |
| post-LLIN | 2010 | Madang | 2094 | 18.6% | 1.2 | 19.5% | 265 | 233 | 8.9 | 2.8 | 6.9 | 2.2 | 0.73 | 0.13 |
|  | 2014 | Madang | 2517 | 9.0% | 1.2 | 17.7% | 93 | 72 | 6.8 | 2.0 | 6.1 | 1.7 | 0.71 | 0.10 |
| all pre-LLIN studies | ≤2006 |  |  | 38.4% | 1.8 | 45.4% |  |  | 9.6 | 0.6 | 7.7 | 0.4 | 0.76 | 0.02 |
| all post-LLIN studies | >2006 |  |  | 12.0% | 1.4 | 26.6% |  |  | 7.7 | 0.5 | 6.5 | 0.4 | 0.71 | 0.02 |
| Overall *P. falciparum* |  |  |  | 25.2% | 1.6 | 35.5% |  |  | 8.7 | 0.4 | 7.1 | 0.3 | 0.74 | 0.02 |
| *P. vivax* |  |  |  |  |  |  |  |  |  |  |  |  |  |  |
| pre-LLIN | 2005 | East Sepik 1 | 1077 | 15.3% | 2.0 | 57.8% | 65 | 64 | 11.5 | 4.2 | 10.9 | 3.9 | 0.83 | 0.09 |
|  | 2005 | East Sepik 2 | 2527 | 35.7% | 2.2 | 58.2% | 187 | 179 | 13.9 | 3.5 | 11.4 | 3.2 | 0.83 | 0.09 |
| post-LLIN | 2012-13 | East Sepik | 2486 | 5.6% | 1.4 | 31.1% | 62 | 47 | 9.3 | 3.1 | 9.3 | 3.0 | 0.80 | 0.08 |

N – number of samples collected per study, MOI – multiplicity of infection, n<sub>s</sub> – number of samples selected for microsatellite genotyping, n<sub>g</sub> – number of samples successfully genotyped, A - the number of alleles, SD- standard deviation, He - expected heterozygosity, Rs - allelic richness, \*N and prevalence (by LDR-FMA for ESP 1 and ESP 2 and qPCR for all other datasets) from [34, 49, 50, 57, 58, 60, 61], \*\* MOI and proportion multiclone infections from *Pfmsp2* or *Pvmsp1-f3* and MS2 and/or MS16 genotyping from [34, 49, 50, 58].

**Table S2.** Estimates of multi-locus linkage disequilibrium single vs all haplotypes.

| all infections |  |  |  | single infections |  |  |
| --- | --- | --- | --- | --- | --- | --- |
| *P. falciparum* | n | $I_A^S$ | p-value | n | $I_A^S$ | p-value |
| East Sepik I 2005 | 15 | -0.0038 | 0.566 | 38 | -0.0076* | 0.718 |
| East Sepik II 2005 | 103 | 0.0004 | 0.446 | 96 | -0.0008* | 0.561 |
| East Sepik 2012-13 | 21 | 0.0454 | **0.002** | 15 | 0.0436* | **0.016** |
| Madang 2006 | 81 | 0.0108 | **0.011** |  |  |  |
| Malala | 17 | -0.0085 | 0.702 | 22 | 0.0074 | 0.708 |
| Mugil | 20 | 0.008 | 0.273 | 21 | 0.0044 | 0.377 |
| Utu | 44 | 0.003 | 0.347 | 36 | 0.0033 | 0.352 |
| Madang 2010 | 82 | 0.0113 | **0.021** | 54 | 0.0211 | **0.00391** |
| Madang 2014 | 33 | 0.0725† | **<0.00001** | 33 | 0.0725 | **<0.00001** |
| *P. vivax* | n | $I_A^S$ | p-value | n | $I_A^S$ | p-value |
| East Sepik I 2005 | 48 | 0.0066 | 0.099 | 26 | 0.0107 | 0.13 |
| East Sepik II 2005 | 37 | 0.0064 | 0.149 | 6 | -0.0085 | 0.69 |
| East Sepik 2012-13 | 20 | 0.2154 | **<0.00001** | 11 | 0.192 | **<0.00001** |
| Madang 2006 | 94 | -0.0022 | 0.796 | 37 | -0.0024 | 0.635 |
| Madang 2010 | 80 | 0.0113 | **0.0005** | 60 | 0.0203 | **<0.00001** |
| Madang 2014 | 24 | 0.0073 | 0.209 | 20 | 0.0100 | 0.184 |

\* data from Schultz et al. 2010
† All isolates single clone infections

**Table S3:**
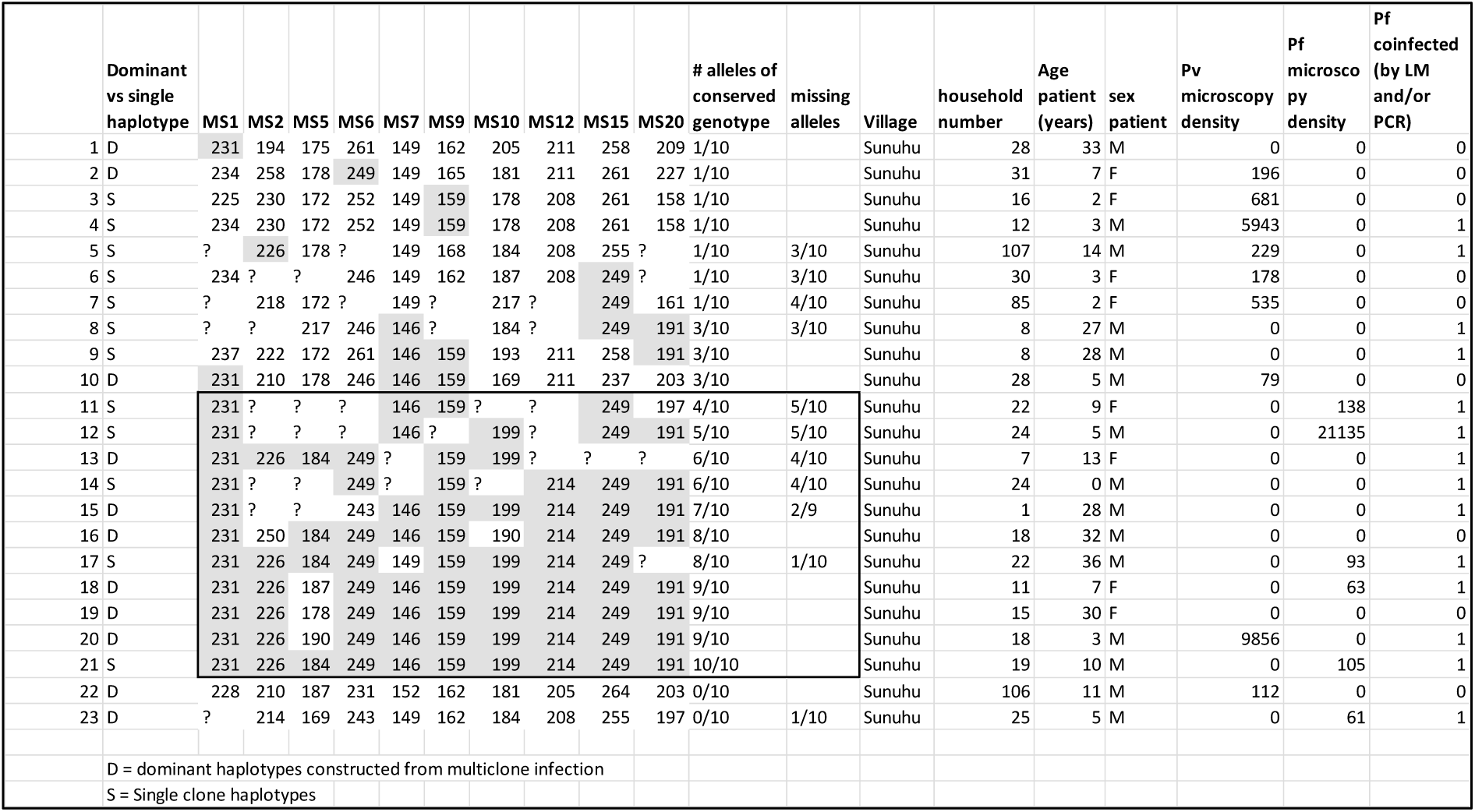
Observed *P. vivax* haplotypes in Sunuhu, ESP 2012-13. Shared alleles from the near-clonal haplotype are shaded grey.

**Table S4:** Estimates of multi-locus linkage disequilibrium (mLD) for *P. falciparum* sub-populations in Madang province.

|  |  | all infections |  |  | unique infections |  |  |
| --- | --- | --- | --- | --- | --- | --- | --- |
| ***P. falciparum*** |  | **n** | **$I_A^S$** | **p-value** | **n** | **$I_A^S$** | **p-value** |
| Malala | 2006 | 17 | -0.0085 | 0.702 | 17 | -0.0085 | 0.702 |
| Mugil | 2006 | 20 | 0.0080 | 0.273 | 20 | 0.0080 | 0.273 |
| Utu | 2006 | 44 | 0.0030 | 0.347 | 44 | 0.0030 | 0.347 |
| Malala | 2010 | 43 | 0.0203 | **0.0118** | 37 | -0.0092 | 0.863 |
| Mugil | 2010 | 33 | 0.0381 | **0.00045** | 29 | 0.0108 | 0.139 |
| Utu | 2010 | 6 | 0.0206 | 0.313 | 6 | 0.0206 | 0.313 |
| Malala | 2014 | 24 | 0.1090 | **<0.00001** | 17 | 0.0160 | 0.171 |
| Mugil | 2014 | 6 | 0.4442 | **<0.00001** | 5 | 0.2810 | **0.00116** |
| Utu | 2014 | 3 | 0.0625 | 0.555 | 3 | 0.0625 | 0.555 |

**Figure S1.**
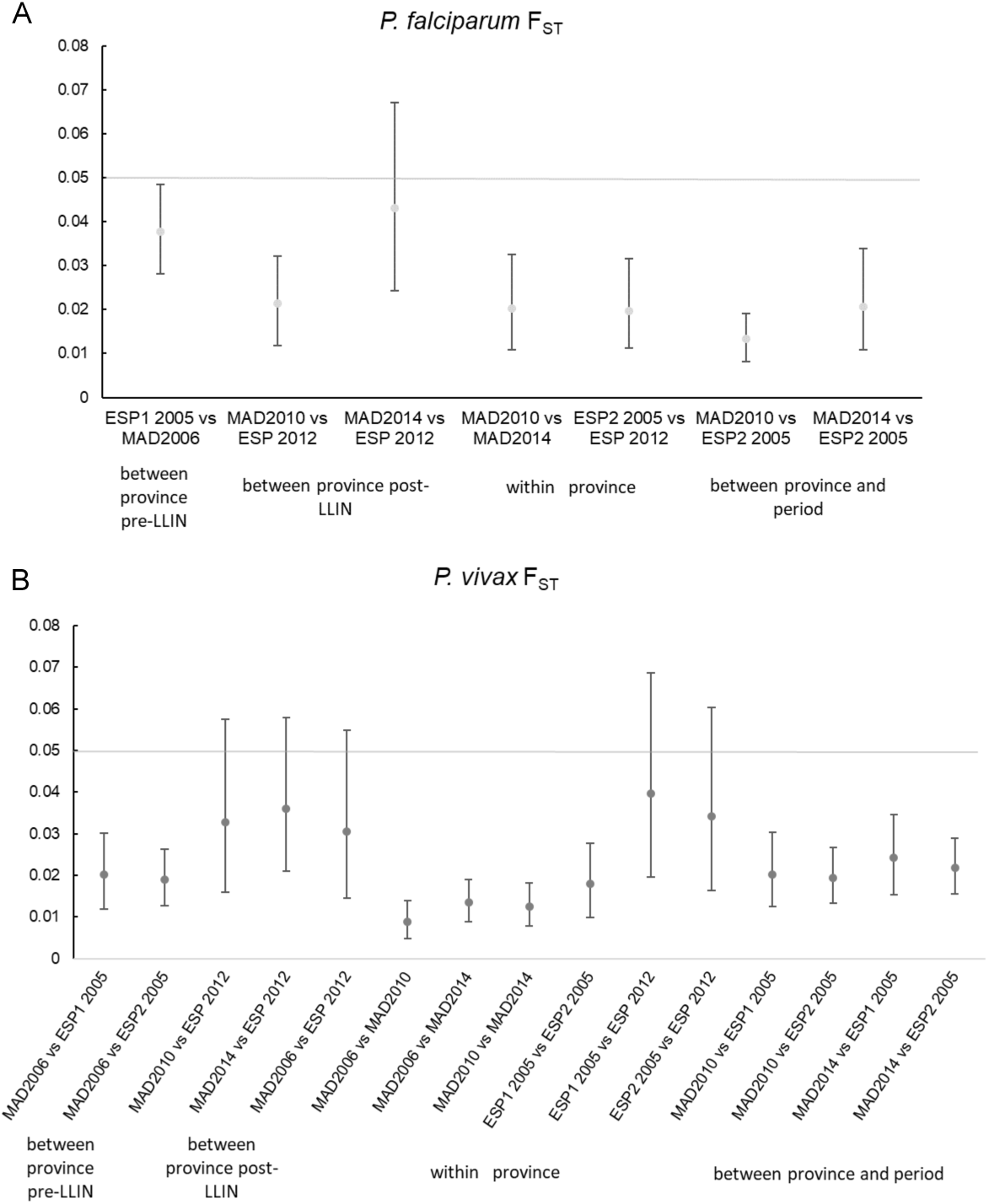
Estimates of genetic differentiation among *P. falciparum* and *P. vivax* populations pre- and post-LLIN mass-distributions. Pairwise F_ST_ values (Weir and Cockerham) for A) *P. falciparum* and B) *P. vivax*. Pairwise F_ST_ values and 95% confidence intervals were estimated with 1000 bootstraps using the diveRsity package in R.

**Figure S2.**
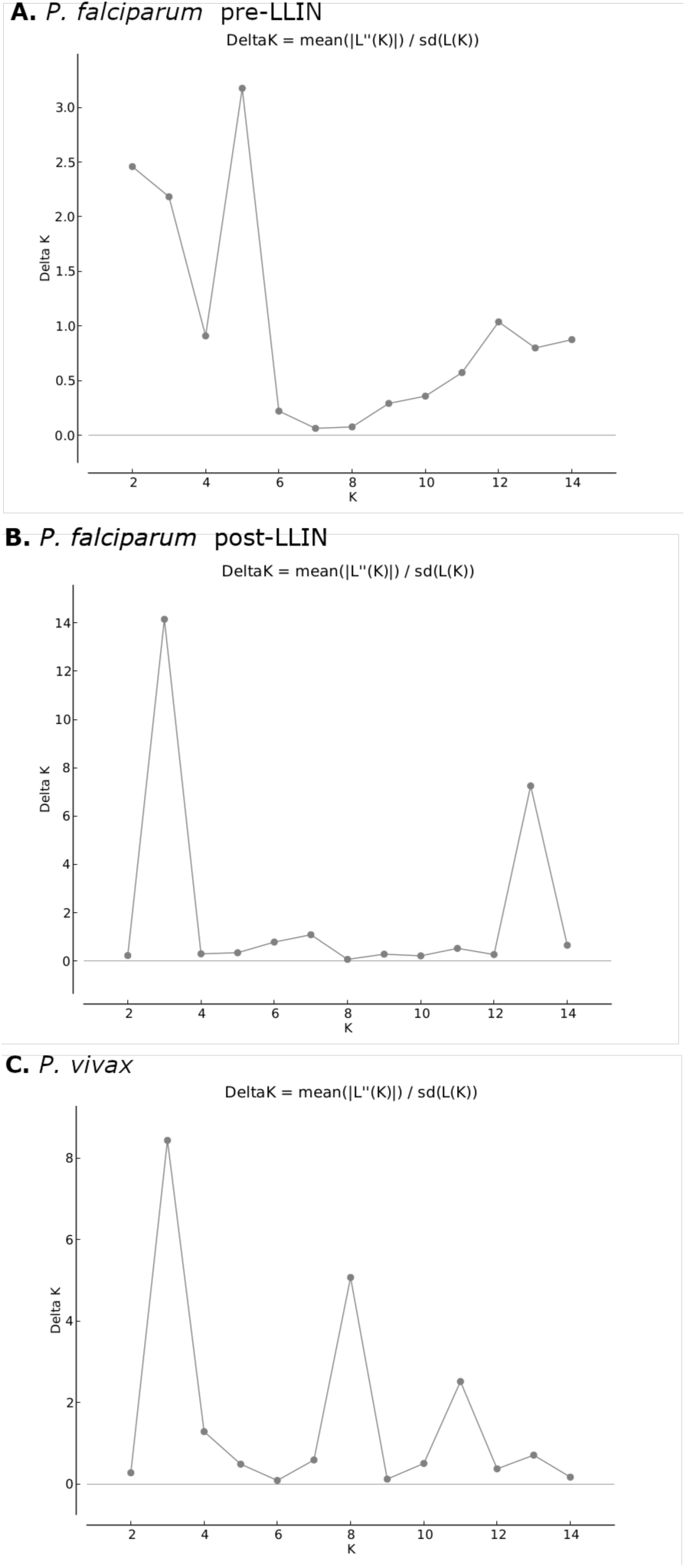
Delta K was calculated to determine the most likely K (ie. number of populations) for the admixture plot for *P. falciparum* pre-LLIN (A) and post-LLIN (B) populations and *P. vivax* populations (C).

